# EMDB - the Electron Microscopy Data Bank

**DOI:** 10.1101/2023.10.03.560672

**Authors:** Jack Turner, The wwPDB Consortium

**Affiliations:** Cellular Structure and 3D Bioimaging, European Molecular Biology Laboratory, European Bioinformatics Institute (EMBL-EBI), Wellcome Genome Campus, Hinxton, Cambridgeshire, CB10 1SD, United Kingdom

## Abstract

The Electron Microscopy Data Bank (EMDB) is the archive of three-dimensional electron microscopy (3DEM) maps of biological specimens. As of 2021, EMDB has been managed by the Worldwide Protein Data Bank (wwPDB) as a wwPDB Core Archive. Today, the EMDB houses over 29,000 entries with maps containing cells, organelles, viruses, complexes and macromolecules. Herein, we provide an overview of the rapidly growing EMDB archive, including its current holdings, recent updates, and future plans.

## Introduction

Cryogenic-sample Electron Microscopy and Tomography (cryo–EM and cryo-ET respectively) (1), together referred to as 3DEM, now represent leading structure determination techniques in the field of structural biology (2–4). Since the advent of the “Resolution Revolution” (5) 3DEM is routinely being used to study structures on scales from atoms (6) to organisms (EMPIAR-11275, to be published). Recent advances also highlight the technique’s applicability to the study of structurally and conformationally heterogeneous complexes (7–10).

In 2002, the Macromolecular Structure Database group, now known as the Protein Data Bank in Europe (PDBe) (11), established the Electron Microscopy Data Bank (EMDB) for the archiving and dissemination of 3DEM volumes (12). The EMDB is headquartered at the European Molecular Biology Laboratory’s European Bioinformatics Institute (EMBL-EBI) in Hinxton, UK. In 2007, the collaboration expanded to include the Research Collaboratory for Structural Bioinformatics (RCSB) (13) and the National Center for Macromolecular Imaging (NCMI) (14). Protein Data Bank in Japan (PDBj) (15) joined the efforts in 2013. In 2021, the EMDB archive became a core wwPDB archive alongside the PDB (16) and BMRB (17), with the EMDB team at EMBL-EBI serving as the wwPDB-designated Archive Keeper for the EMDB core archive. Protein Data Bank China (PDBc) recently joined the wwPDB as an associate member (18), further expanding the international collaboration managing these archives. wwPDB core archives are made available at no charge and with no limitation on usage under the CC0 1.0 Creative Commons licence (https://creativecommons.org/share-your-work/public-domain/cc0/). All three core archives managed jointly by the wwPDB operate under the FAIR principles of Findability, Accessibility, Interoperability, and Reusability (19). Following the FAIR principles facilitates data discovery, collaboration, and data reproducibility - all of which are important for accelerating research and innovation.

Interoperation between archives is essential for serving the complete data associated with a 3DEM experiment. The 3DEM Coulomb potential map (henceforth referred to as ‘map’ or ‘volume’) is stored in the EMDB archive, whilst the derived coordinate models are held in the PDB archive. The raw data from which the map is derived are collected by the Electron Microscopy Public Image Archive (EMPIAR) (20). In addition to the aforementioned structural archives, entries in the EMDB can also be associated with entries in the AlphaFold Protein Structure Database (21) and Small Angle Scattering Biological Data Bank (SASBDB) (22).

wwPDB members support its core mission of sustaining freely accessible, interoperating Core Archives of structure data and metadata for biological macromolecules as an enduring public good to promote basic and applied research and education across the sciences (16). Herein, we provide an overview of the recent developments in the EMDB archive including its content, deposition protocols, use, and future prospects.

## Archive Content

### Growth and Statistics

The EMDB archives structures from the methods single particle analysis (SPA), subtomogram average (STA), helical reconstruction (HR), tomography, and electron crystallography (EC). All of these techniques produce maps with the exception of EC, which produces diffraction data from which maps can be generated.

EMDB holds > 29,000 entries as of 31st August 2023, approximately 55% of which have associated atomic coordinates housed in the PDB (**Figure 1a**). The number of EMDB entries released per year is growing exponentially (**Figure 1b)**. In comparison, the growth of X-ray data in the PDB has remained steady since 2016 at around 10,000 entries per year. Based on these trends, it is expected that the number of EM entries released per year will surpass X-ray releases by the end of 2024. Data available at the end of August 2023 suggest ∼7785 EMDB entries (in agreement with the trendline (**Figure 1b)**) and ∼9272 X-ray entries (slightly short of the total predicted by the trendline) will be released in 2023 (this prediction assumes consistent release numbers throughout the year). Whilst exponential growth is unlikely to continue indefinitely, at the current rate EMDB is predicted to hold 50,000 entries in 2025 and 100,000 entries in 2028 (a doubling time of ∼2.5 years), with ∼13,500 and ∼31,500 releases predicted in the same years.

**Figure 1:**
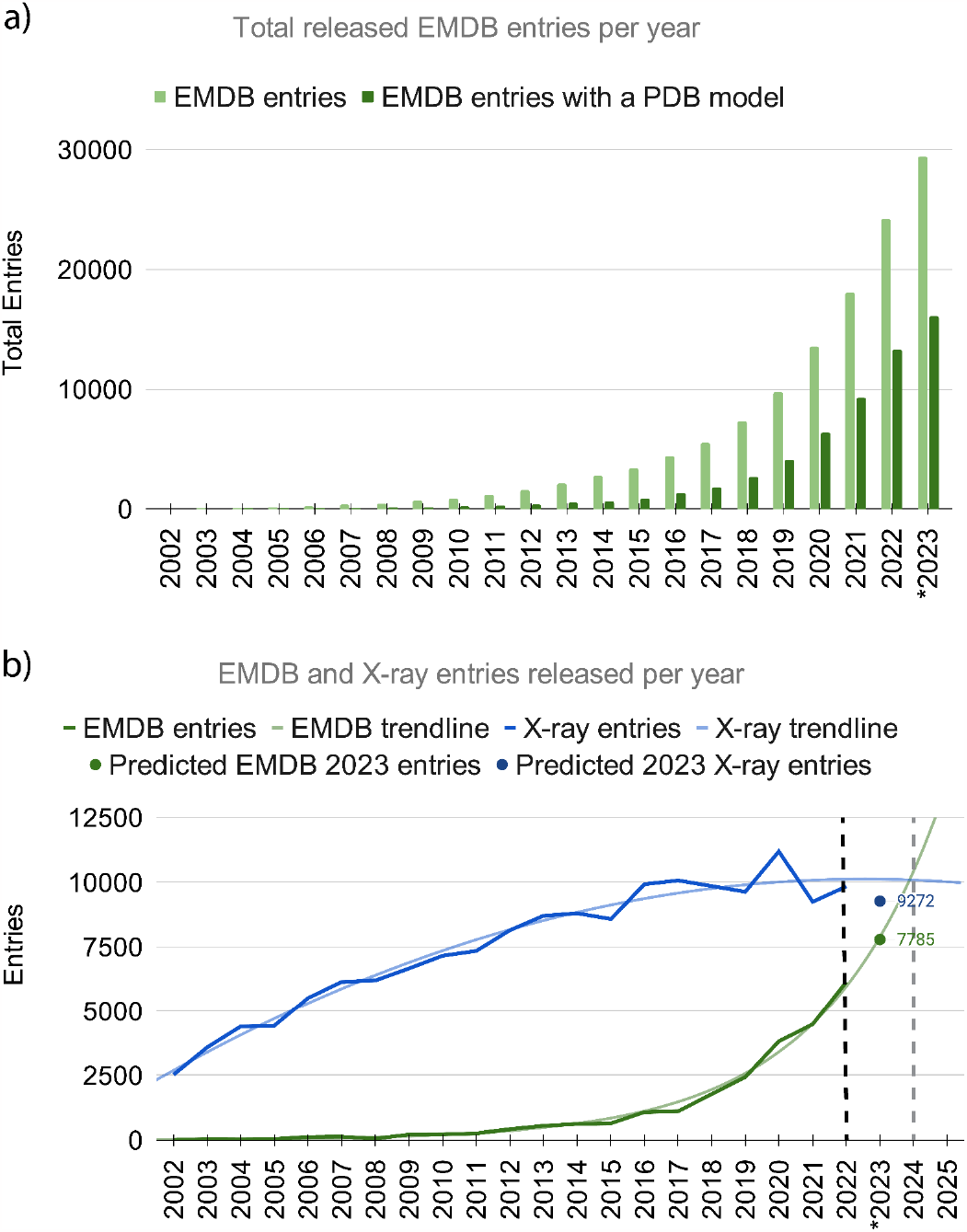
**(a)** Total number of EMDB entries and total number with a coordinate model in the Protein Data Bank released per year. * 2023 data are through August 31st. **(b)** The number of entries released to the EMDB per year compared with the number of X-ray entries released per year. Data available up until 2022 (black dashed line) after which trendlines are plotted to show the predicted crossover point (grey dashed line, 2024). A polynomial line (x^2^ with an R^2^ = 0.97) was fitted to the X-ray data and an exponential curve (R^2^ = 0.99) was fitted to the EMDB data. * predicted numbers according to (n/8)*12 where n is the number of entries released as of August 31st.

The proportion of entries in the EMDB archive that corresponds to each methodology is shown in **Figure 2a**. SPA is by far the most popular method deposited to the archive, making up 82.8% of the total archive at the end of 2022, an increase of > 4% since 2016 (23). STA, Tomography, HR, and EC make up the remaining 17.2% in descending abundance. Using metadata harvested from the EMDB, the effect of the Resolution Revolution (5) can be visualised (**Figure 2b)**. A rapid improvement in the average resolution of released structures has been seen since 2014. Resolutions better than 4 Å have made up > 50% of releases since 2019 (1131 entries in 2019 rising to 4033 in 2022), and sub-3 Å structures make up almost 25% of 2022’s released entries (216 entries in 2019 rising to 1299 in 2022). Higher resolution 3DEM volumes enable more accurate building of atomic coordinate models. The highest resolution single particle-based structure in the archive is currently that of an apoferritin protein at ∼1.15 Å resolution (EMD-11668, 7A6A (6)). Diffraction methods push the resolution even further with structures achieving resolutions of ∼0.6 Å (*e*.*g*., EMD-0698, 6KJ3 (24)).

**Figure 2:**
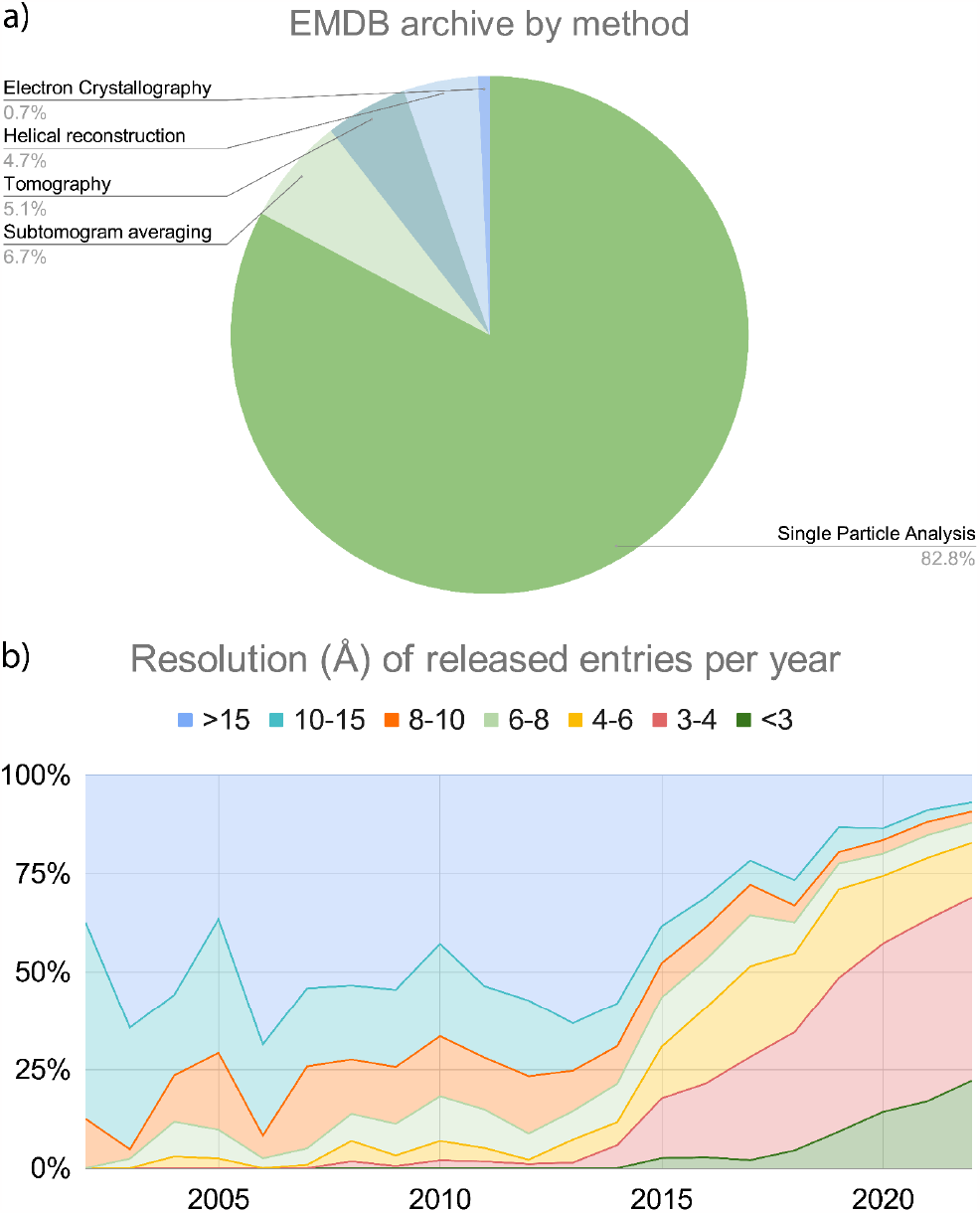
**(a)** Proportion of the EMDB archive made up by each accepted method from 2002 - 2022. **(b)** Proportion of releases in various resolution (Å) shells per year from 2002 - 2022.

### Archive Data

Each EMDB entry describes a 3DEM experiment. All entries contain a primary 3DEM volume around which an entry is based, except EC entries, which must be deposited with structure factors and may optionally include a primary map. Unfiltered, unmasked, and unsharpened half-maps, generated as standard in most SPA, STA, and HR workflows, are present for all relevant methodologies deposited after February 25th, 2022 (https://www.wwpdb.org/news/news?year=2022#6218da3152988f064bf8c4a3). Each entry is presented with an image file provided by the depositor, enabling them to showcase the primary map according to their preferences. A number of optional, additional files may also be present in an entry, including additional maps, masks, an Fourier Shell Correlation (FSC) curve, and layer line files. **Table 1** gives an overview of all files presented in an entry.

**Table 1:** Summary of an entry’s contents in the archive.

| Category (number) | Description |
| --- | --- |
| Primary map (1) | EM map or tomogram that is shown and described in the associated publication |
| Half-maps (0 or 2) | Unfiltered, unsharpened, and unmasked raw half-maps for single-particle analysis, single-particle-based helical reconstructions, or subtomogram averaging |
| Masks (0 or more) | Primary/raw map masks, segmentation/focused refinement masks, and half-map masks |
| Additional cryo-EM maps (0 or more) | Examples include difference maps, maps showing alternative conformations and/or compositions, and maps with different processing (for example filtering, sharpening, and masking) |
| Auxiliary files (0 or more) | Examples include author-determined FSC curves (half-maps, map-model, ...), structure factors, and layer lines |
| XML metadata files (1) | XML file containing an entry's metadata |
| Validation Report (3) | Map-only validation reports (3 file types), map and model summary report, and map and model full report. |

An entry’s primary volume must be adequately described with additional metadata to ensure adherence to FAIR principles (19). These metadata are stored in Extensible Markup Language (XML) with items and attributes defined in the EMDB data model (www.ebi.ac.uk/emdb/documentation#version30). Utilised definitions include a hierarchy of information which allows description of cellular structures, supramolecules, and macromolecules, all of which may exist in a single map. Various other experimental metadata are also present in these data files, including information on specimen preparation, microscopy instrumentation, data collection protocol, and software used during map generation and processing. The EMDB data model includes definitions of mandatory items and support for enumerations and allowed data ranges. All metadata files in the EMDB archive can, therefore, be validated against the XML schema.

## Global Data Deposition

Depositions to the EMDB are managed by the wwPDB global OneDep deposition (25), validation (26–28), and biocuration software system (29). OneDep is a unified deposition system for 3DEM, X-ray, and NMR data. OneDep is hosted at wwPDB data centres around the world (located in the USA, UK, and Japan), providing a consistent deposition experience regardless of a structural biologist’s geographic location. Biocuration of incoming entries is geographically distributed among wwPDB partner sites as follows: RCSB PDB handles all depositions from the Americas and Oceania, PDBe and the EMDB team at EMBL-EBI manage the depositions from Europe and Africa, and PDBj and PDBc handle all depositions from Asia and the Middle East. This arrangement divides the effort of biocuration resources and ensures that depositors can communicate with wwPDB biocurators within, or close to, their local timezone.

The OneDep system makes use of the PDBx/mmCIF framework during deposition, validation, and biocuration. All data and metadata related to a deposition are defined in the EMDB data model, which in turn informs the PDBx/mmCIF dictionary (mmcif.wwpdb.org) (30). Thus for 3DEM, the PDBx/mmcif dictionary faithfully represents the EMDB model described in the EMDB XML schema. Data specific to 3DEM methods in the PDBx/mmCIF dictionary uses the “em” namespace (*e*.*g*., ‘em_imaging’ for metadata on the electron microscope setup) for which a full list of categories, broken down by groups, can be found here: https://mmcif.wwpdb.org/dictionaries/mmcif_em.dic/Groups/index.html. The dictionary definitions can also include rules such as relationships between different data items, enumerations, and allowed ranges. Finally, the PDBx/mmCIF format is extensible, allowing the dictionary to grow with the data models for all three wwPDB core archives (PDB, EMDB, and BMRB).

The OneDep deposition interface provides a set of comprehensive web forms for capturing metadata. Cross-checking is automatically carried out at various stages to ensure the validity of entered data. If metadata is provided within the uploaded mmCIF file, it is used to automatically populate the relevant items in the deposition interface. Functionality was recently added to allow the deposition of a metadata mmCIF file when atomic coordinates are not being deposited, which allows the deposition interface to be automatically filled for map-only 3DEM depositions. Biocuration involves various steps including ligand and sequence annotation, validation, and data integrity checks.

At the end of the deposition, validation, and biocuration processes a wwPDB validation report is provided to the depositor. This report contains a range of community-recommended validation metrics for both the map and, if present, the atomic coordinates (26, 28, 31). It is strongly recommended by the wwPDB that depositors provide their confidential wwPDB validation reports in PDF format to scientific journals when submitting related manuscripts (16). Validation metrics related to 3DEM maps are implemented in a tiered approach, allowing testing and gathering of community feedback by presenting the metrics on the EMDB website (tiers 1 and 2) before implementation into the wwPDB validation report (tier 3). Q-score (32) is the most recent metric to be added to tier 3, providing a new map-model fit metric which complements the previously implemented atom-inclusion score.

Upon depositor request, manuscript publication (including online preprint servers), or if one year has elapsed since the time of submission, an entry is staged for public release in the next weekly EMDB update cycle. Entries can be staged for release from Monday to Thursday in any given week. Release of the atomic coordinates from the PDB cannot proceed without release of the experimental data, and likewise release of atomic coordinates cannot precede release of the associated EMDB entry. Every Thursday, after all entries are staged, the EMDB archive begins the release process, which includes data integrity checks, running of automated validation and data-enrichment pipelines, and manual inspection of validation outputs for all entries. Depositors are contacted in the event of peculiarities noticed during final validation checks. New data are then made public every Wednesday at 00:00 UTC.

## Data Dissemination

The EMDB archive is served *via* an FTP (https://ftp.ebi.ac.uk/pub/databases/emdb/), which is mirrored by wwPDB (https://files.wwpdb.org/) and PDBj (https://files.pdbj.org/pub/emdb/). The EMDB FTP provides access to map, image, FSC (if available), metadata, and validation files for released and obsoleted entries. The EMDB FTP also serves files describing the EMDB data model and the status of all released and unreleased 3DEM entries. Download and unique download request statistics from the EMDB FTP are shown in **Figure 3a**. In 2022, 6,597 unique IP addresses requested downloads. The total volume of data downloaded exceeded 24 TB (Total archive size at end of 2022 was 8.9 TB), an ∼8TB increase from the previous year. When assessing total downloads of EMDB data across all wwPDB FTPs (**Figure 3b**) seasonal variation is observed. Most download requests occurred during Q4 of 2020 and 2021, in 2022 download requests peaked slightly earlier, in Q3.

**Figure 3:**
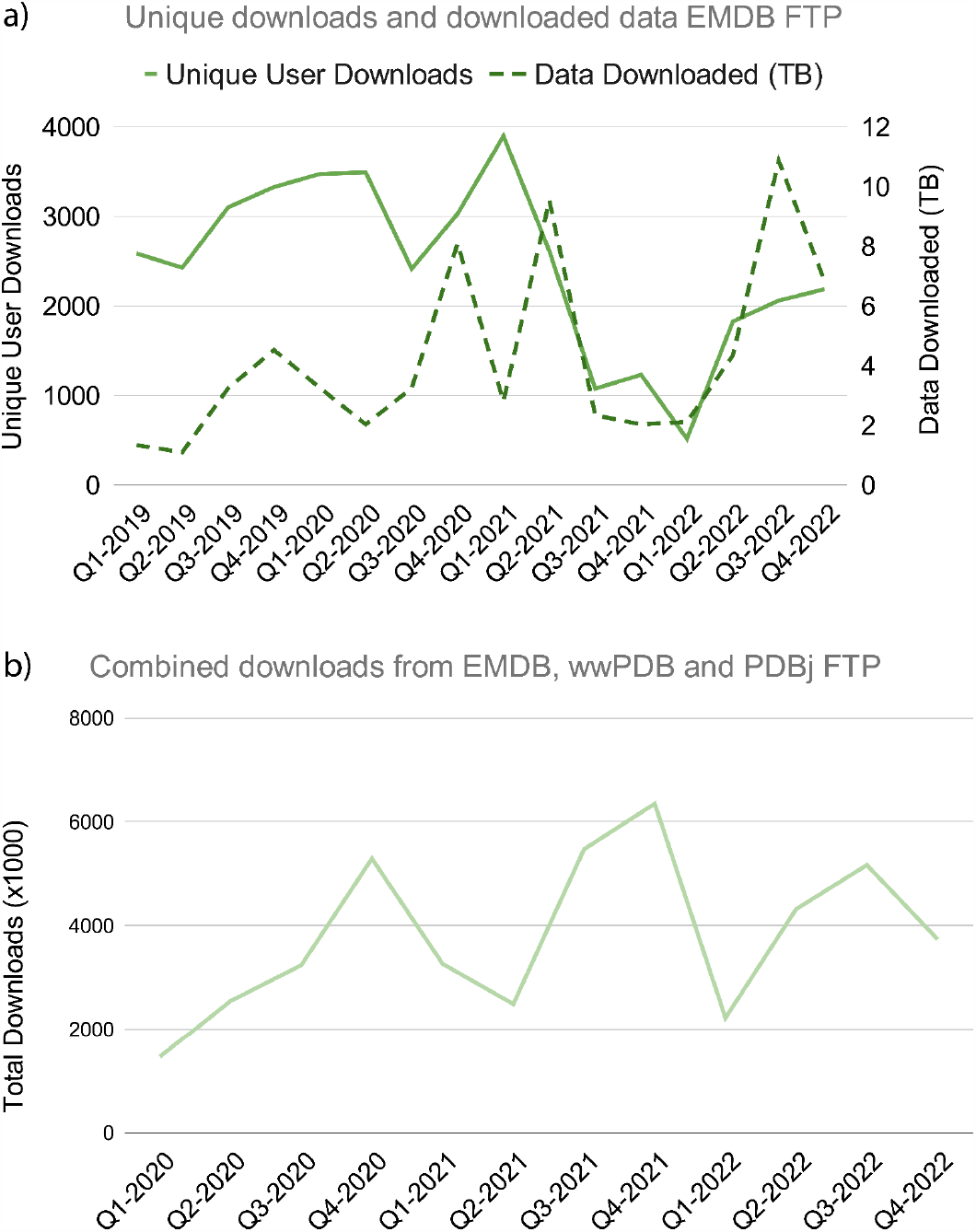
**(a)** Quarterly statistics on the number of download requests made by unique IP addresses (solid line) and the total quantity of data served (dashed line). **(b)** Quarterly statistics on the total number of download requests for EMDB data from all wwPDB FTPs.

Data, news, and statistics relevant to the EMDB can be found on the EMDB website (www.ebi.ac.uk/emdb/). News and statistics pertaining to the wider wwPDB, including the EMDB, are available on the wwPDB website (www.wwpdb.org). Volume, image, and any additional data are served in map format (ftp.ebi.ac.uk/pub/databases/emdb/doc/Map-format/). Metadata is provided in XML format, with the data model described in the ‘docs/’ section of the FTP.

## Archive Updates

A major change to the EMDB archive involved the transition to a new version of the EMDB data model, which was completed in 2021. The new data model contains a rich set of metadata and supports hierarchical description of the sample composition. Descriptions of both molecular and cellular samples are supported, and description of the overall EM experiment has improved relative to the previous version of the data model, which was originally developed for use with EMDep (33), the previous 3DEM data deposition system. With this change 3DEM experiments can be more appropriately described.

When extending the EMDB data model, archive remediation is required in order to provide the new information for historical entries, where possible. For example, a recent addition to the data model is the functionality to relate entries to the SASBDB. These data are now available for newly released entries and a remediation will be carried out to serve this information for all historical entries. When major changes are made to the archive and/or policies, they are communicated via the EMDB website and wwPDB channels, including emails to major mailing lists (*e*.*g*. CCPEM and 3DEM).

## Community Engagement

### Expert User Groups

Community engagement is at the heart of archive management. To this end, the wwPDB is guided by an expert international advisory board. The board advises on the activities of the wwPDB partners annually; outcomes of the annual board meetings are published on the wwPDB website (http://www.wwpdb.org/about/advisory).

In addition to the yearly advisory board several workshops specific to the planning of EMDB’s future endeavours have been conducted. Community experts were engaged for the purposes of ensuring optimal data management (34), annotation of cellular data (35), and ensuring integration of data between multiple resources (36). The validation methods in electron microscopy are constantly evolving, and no one metric can provide an overall description of volume quality. An expert Validation Task Force was set up and first met in 2010 (31), with a second meeting held in 2020. This group contributes and advises on the validation metrics EMDB includes in its Validation Analysis resource (28).

### Training and Outreach

Training is essential to ensure that the community has an understanding and awareness of the full suite of wwPDB archives and the resources they offer. EMDB staff have provided training globally at various cryo-EM courses, including the Cold Spring Harbour Laboratory cryo-EM course; the Sao Paulo Cryo-EM course; and the EMBL-EBI bioinformatics courses. In addition, EMDB staff regularly present posters at relevant conferences including CCPEM, Tomography Congress 2022, and the 3DEM GRC, giving them the chance to engage and gather feedback from the community.

The EMDB and EMPIAR teams collaborate to run an X (previously known as Twitter) account and YouTube channel, both with the handle @EMDB_EMPIAR. Several tweets are posted to the X account per week with content ranging from interesting structures from the week’s release to advertisements for team job opportunities. The EMDB YouTube channel hosts various videos including recorded talks from meetings and tutorials on how to make the most of EMDB website features.

## Current Trends and Future Outlook

### Rapid Growth

The EMDB archive continues to experience exponential growth year-on-year (**Figure 1**), while at the same time depositors are generating an increased quantity of entries per publication (**Figure 4**). wwPDB partners are addressing these developments through improvements to the OneDep deposition-validation-biocuration system. Automation of the biocuration and validation workflows is being constantly reviewed, planned, and implemented. Depositor experience and efficiency also represent crucial considerations. The new ORCID login feature allows depositors to view all of their entries within the browser, rather than having to manually curate their own logbook of deposition IDs. A deposition API is also in development, this will allow software packages to directly deposit data into the OneDep system, in the most favourable cases entirely obviating the need for depositor intervention.

**Figure 4:**
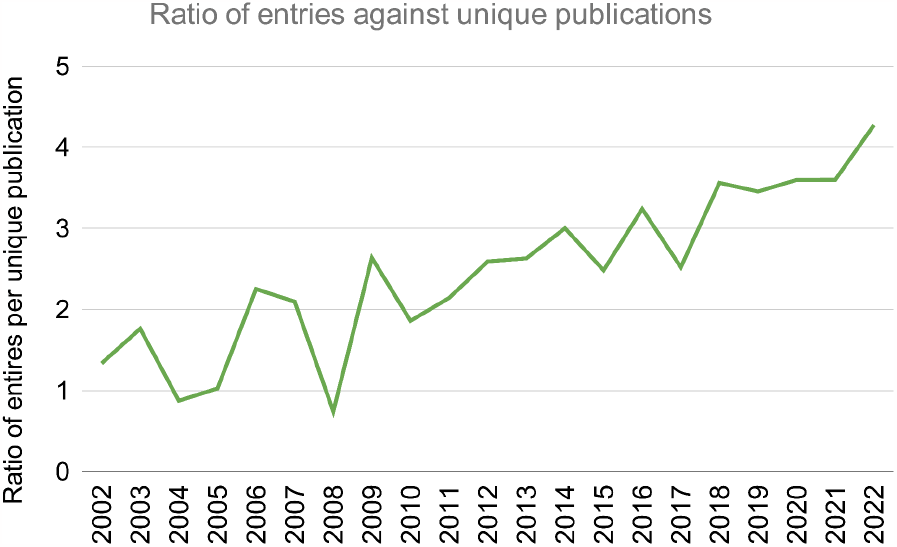
Ratio of entries per unique publication plotted per year.

### Rapidly Evolving Science

Every structure determination submethod supported by EMDB is witnessing continuous innovation and improvement. Single particle-based methods (SPA and STA) can now be used to analyse conformational variability, sometimes referred to as heterogeneity analysis (7–10). This approach allows multiple structural states to be resolved from a single imaging dataset. Simultaneously, tomography is seeing explosive growth in the amount of data that can be recorded per unit time (37, 38). Such advances require frequent updates to the EMDB data model and PDBx/mmCIF dictionary. The EMDB plans to establish a wwPDB 3DEM working group to accelerate implementation of such additions and promote adoption of data and methodological standards across the community.

Software development in the field of artificial intelligence/machine learning (AI/ML) is also seeing rapid growth. EMDB data have already been used in the development of new AI-based software tools for various applications, including particle picking (39), particle pruning (40), map sharpening (41), local resolution estimation (42), secondary structure detection (43), and residue-level quality estimation (44). The EMDB endeavours to continue supporting the development of new ML algorithms through expert curation, and thorough and accurate labelling of entries *via* the EMDB data model.

Programmatic collection and deposition of data is one avenue for improving metadata accuracy and completeness. The wwPDB is currently working to implement programmatic deposition of entries facilitated by a deposition API. The wwPDB is currently working with developers from popular structural biology software packages to accelerate the adoption of the deposition API.

### Looking Forward

3DEM enables structural understanding of biomolecules, cells, tissues, organs, and organisms. As the field continues to expand and evolve, the importance of archiving and appropriately describing new types of data will become ever more important. To this end, the EMDB plans to continue to expand the 3DEM data model in collaboration with the to-be-established wwPDB 3DEM and wwPDB PDBx/mmCIF working groups. Examples of imminent enhancements include provenance description of synthesised macromolecules and appropriate labelling of composite maps. Planned improvement of metadata describing 3DEM experiments will both support and enhance interoperability of archived data. This process will include archiving of metadata in mmCIF format at EMDB, in addition to the already available XML format.

The EMDB Validation Analysis software package (28) generates an extensive set of validation metrics, which enable assessment of various map/volume and map-model features. Comprehensive metrics are important, but the sheer volume of metrics can be overwhelming to some users, particularly those less familiar with 3DEM techniques. Furthermore, calculating an ever-expanding set of metrics for every entry in the exponentially growing archive will represent a significant computational burden. Future thought will need to be aimed at summarising important results from the Validation Analysis pipeline, whilst limiting the computational and environmental costs of running the software.

Summarising results is likely to take a similar form to the sliders already presented within wwPDB validation reports for atomic models deposited to the PDB archive.

Finally, validation of deposited metadata is essential for ensuring that the archive retains its value in the longer term. The wwPDB has recently implemented a number of 3DEM-related checks within the OneDep system to improve reliability. For example, minimum defocus values can no longer be greater than maximum defocus values and provided pixel sizes of volumes are compared with those reported in the volume header. Additionally, deposition of atomic coordinates that include parts of structures extending beyond the volume bounding box is now stopped at the file upload stage. More inter-file checks are in development, including assessment of whether uploaded half-maps are identical, or whether the primary map and half-maps are offset from one another. In rare cases, depositors accidentally select the wrong methodology at the beginning of the deposition process. To mitigate this issue the EMDB is experimenting with deep learning approaches to predict the experimental method from which the volume was derived. Taken together, these checks will help to ensure that data archived within the EMDB are both as accurate as possible and adhere to the FAIR principles.

## Current wwPDB Consortium Members with Affiliations

EMDB

Jack Turner^1^, Sanja Abbott^1^, Neli Fonseca^1^, Lucas Carrijo^1^, Amudha Kumari Duraisamy^1^, Osman Salih^1^, Zhe Wang^1^, Gerard Kleywegt^1^, Kyle L. Morris^1^, Ardan Patwardhan^1^

RCSB PDB

Stephen K. Burley^2,3,4,5^, Gregg Crichlow^2^, Zukang Feng^2^, Justin W. Flatt^2^, Sutapa Ghosh^2^, Brian P. Hudson^2^, Catherine L. Lawson^2^, Yuhe Liang^2^, Ezra Peisach^2^, Irina Persikova^2^, Monica Sekharan^2^, Chenghua Shao^2^, Jasmine Young^2^

PDBe

Sameer Velankar^6^, David Armstrong^6^, Marcus Bage^6^, Wesley Morellato Bueno^6^, Genevieve Evans^6^, Romana Gaborova^7^, Sudakshina Ganguly^6^, Deepti Gupta^6^, Deborah Harrus^6^, Ahsan Tanweer^6^, Manju Bansal^8^, Vetriselvi Rangannan^8^

PDBj

Genji Kurisu^9,10^, Hasumi Cho^9^, Yasuyo Ikegawa^9^, Yumiko Kengaku^10^, Ju Yaen Kim^10^, Satomi Niwa^10^, Junko Sato^10^, Ayako Takuwa^10^, Jian Yu^10^

BMRB

Jeffrey C. Hoch^11^

PDBc

Wenqing Xu^12,13^, Weizhe Zhang^12^, Xiaodan Ma^12^

1. Cellular Structure and 3D Bioimaging, European Molecular Biology Laboratory, European Bioinformatics Institute (EMBL-EBI), Wellcome Genome Campus, Hinxton, Cambridgeshire, CB10 1SD, United Kingdom
2. Research Collaboratory for Structural Bioinformatics Protein Data Bank, Institute for Quantitative Biomedicine, Rutgers, The State University of New Jersey, Piscataway, NJ 08854, USA
3. Department of Chemistry and Chemical Biology, Rutgers, The State University of New Jersey, Piscataway, NJ 08854, USA
4. Cancer Institute of New Jersey, Rutgers, The State University of New Jersey, New Brunswick, NJ 08901, USA
5. Research Collaboratory for Structural Bioinformatics Protein Data Bank, San Diego Supercomputer Center, University of California, La Jolla, CA 92093, USA
6. Protein Data Bank in Europe, European Molecular Biology Laboratory, European Bioinformatics Institute (EMBL-EBI), Wellcome Genome Campus, Hinxton, Cambridgeshire, CB10 1SD, United Kingdom
7. CEITEC - Central European Institute of Technology, Masaryk University, Kamenice 5, 62500 Brno, Czech Republic
8. Molecular Biophysics Unit, Indian Institute of Science, Bangalore. India.
9. Protein Data Bank Japan, Protein Research Foundation, Minoh, Osaka 562-8686, Japan
10. Protein Data Bank Japan, Institute for Protein Research, Osaka University, Suita, Osaka 565-0871, Japan
11. Biological Magnetic Resonance Data Bank, Department of Molecular Biology and Biophysics, UConn Health, 263 Farmington Ave., Farmington CT 06030-3305 USA
12. National Facility for Protein Science, Shanghai Advanced Research Institute, Chinese Academy of Sciences, Shanghai 201210, China
13. School of Life Science and Technology, ShanghaiTech University, Shanghai 201210, China

